# Rac1 is a downstream effector of PKCα in structural synaptic plasticity

**DOI:** 10.1101/750372

**Authors:** Xun Tu, Ryohei Yasuda, Lesley A Colgan

## Abstract

Structural and functional plasticity of dendritic spines is the basis of animal learning. The calcium-dependent protein kinase C isoform, PKCα, has been suggested to be critical for this actin-dependent plasticity. However, mechanisms linking PKCα and structural plasticity of spines are unknown. Here, we examine the spatiotemporal activation of actin regulators, including small GTPases Rac1, Cdc42 and Ras, in the presence or absence of PKCα during single-spine structural plasticity. Removal of PKCα expression in the postsynapse attenuated Rac1 activation during structural plasticity without affecting Ras or Cdc42 activity. Moreover, disruption of a PDZ binding domain within PKCα led to impaired Rac1 activation and deficits in structural spine remodeling. These results demonstrate that PKCα positively regulates the activation of Rac1 during structural plasticity.

## Introduction

Dendritic spines of pyramidal neurons in the hippocampus undergo activity-dependent structural and functional plasticity that has been reported to be crucial for learning and memory (Lai and Ip, 2013; Lamprecht and LeDoux, 2004; Matsuzaki et al., 2004). This plasticity is mediated by the coordinated regulation of complex signaling networks that transduce short-lived synaptic input into long-lasting biochemical changes to modulate the strength and structure of synapses and, ultimately, animal behavior (Kennedy, 2016; Nishiyama and Yasuda, 2015).

Protein kinase C (PKC) is a family of serine/threonine kinases that have long been implicated as essential for synaptic plasticity, learning and memory (Nelson et al., 2008). Inhibition of PKC blocks LTP induction and also disrupts the maintenance of pre-established LTP (Malinow et al., 1989, 1988). In addition, pharmacologically activating PKC or overexpressing constitutively active PKC potentiates synapses and enhances learning (Hu et al., 1987; Rekart et al., 2005; Zhang et al., 2005). PKC exists as at least 10 isozymes which are categorized into three subfamilies: the classical PKC isozymes (PKCα, PKCβ and PKCγ), the novel isozymes (PKCδ, PKCε, PKCη, and PKCθ) and the atypical isozymes (PKCζ and PKCλ). The classical PKC isozymes, ubiquitously expressed in the brain, transduce signals dependent on Ca^2+^ and diacylglycerol (DAG) (Clark et al., 1991; Ito et al., 1990; Kose et al., 1990; Sossin, 2007). Recently, PKCα was demonstrated to be uniquely required for structural LTP in hippocampal dendritic spines. This specificity was defined by a four amino acid C-terminal PDZ-binding motif (QSAV). Evidence suggests that PKCα activity integrates neurotrophic signaling, including the activation of TrkB, with Ca^2+^ influx through NMDARs to facilitate the induction the plasticity in dendritic spines (Colgan et al., 2018). However, the downstream molecular mechanisms through which PKCα facilitates structural synaptic plasticity remains unknown.

The expression of structural plasticity, through spine enlargement and insertion of additional glutamate receptors, requires actin remodeling through the regulated activity of small GTPases including Rac1, Cdc42 and Ras (Bailey et al., 2015; Kim et al., 2014; Murakoshi et al., 2011; Patterson and Yasuda, 2011). These small GTPases are precisely coordinated across spatiotemporal domains by a complex network of GTPase accelerating proteins (GAPs) and GTPase exchange factors (GEFs), which attenuate or enhance GTPase activation respectively to mediate the cytoskeletal remodeling crucial for activity-dependent spine plasticity (Govek et al., 2005; Lai and Ip, 2013; Saneyoshi et al., 2008).

## Result

Here, we examine whether PKCα regulates small GTPases during the induction of plasticity. In order to study signaling in single spines during structural plasticity, we combined two-photon release of caged glutamate (Matsuzaki et al., 2001), fluorescence resonance energy transfer (FRET)-based sensors, and two-photon fluorescence lifetime imaging microscopy (2pFLIM) (Yasuda, 2012) to monitor the dynamics of intracellular signaling events with high spatiotemporal resolution. Specifically, using previously published FRET sensors, we monitored the spatiotemporal activation of the actin-regulating small GTPases, including Ras (Harvey et al., 2008; Oliveira and Yasuda, 2014; Yasuda et al., 2006), Cdc42 (Murakoshi et al., 2011) and Rac1 (Hedrick et al., 2016) during the induction of structural long-term potentiation (sLTP). Briefly, these sensors are composed of two components: (1) full length GTPase fused to green fluorescent protein (GTPase–eGFP), and (2) a specific GTPase binding domain of a downstream effector fused to two copies of red fluorescent protein (mRFP-effector-mRFP) (Fig. 1a). Activation of the GTPase increases the affinity between the two components of the sensor and the FRET between the fluorophores. Using 2pFLIM we can monitor changes in the binding of the sensor components by quantitatively measuring decreases in the fluorescence lifetime of GFP in order to measure small GTPases activation.

**Figure 1.**
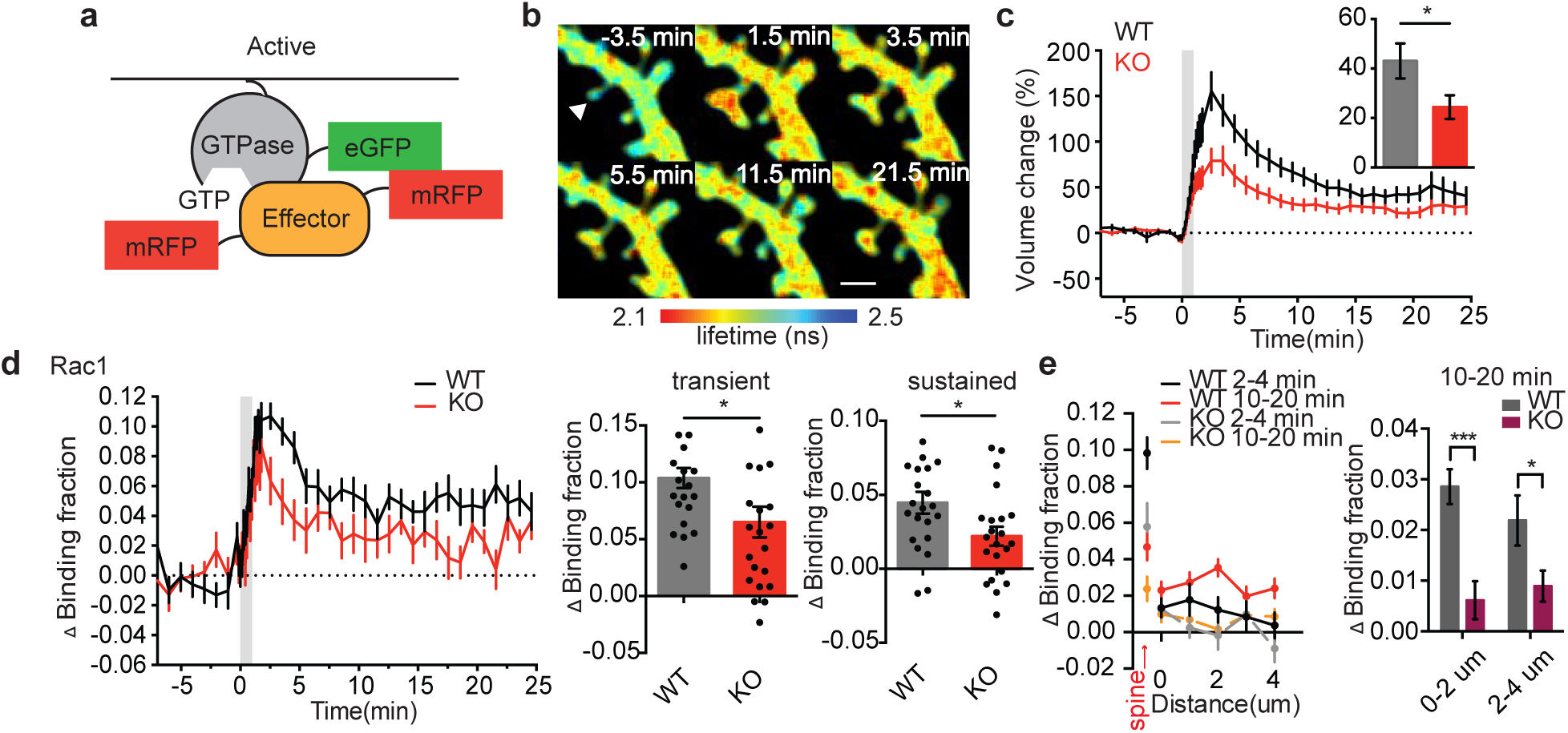
PKC regulates Rac1 activation during sLTP. **a**, Schematic of small GTPase FRET sensors. **b**, 2pFLIM images of Rac1 activation averaged across indicated time points. Arrowhead represents point of uncaging. Warmer colors indicate shorter lifetimes and higher Rac1 activity. Scale bar, 1μm. **c**, Time courses and quantification of sustained (10–25 min) spine volume change induced by glutamate uncaging in neurons expressing Rac1 sensor from WT and KO littermates for PKCα. **d**, Time courses and quantification of transient (1.5-3.5 min) and sustained (10-25 min) Rac1 activation in stimulated spines from WT and PKCα KO littermates. **e**, Spatial profile and quantification of spreading Rac1 activation along the dendrite at indicated times and distances from the stimulated spine in WT and PKCα KO littermates. Data are mean ± s.e.m. Grey shading indicates time of uncaging. *P < 0.05, two-tailed t-test (**c, d**) and two-way ANOVA with Sidak’s mutiple comparisons test (**e**). n (neurons/spines) = 18/22 WT and 19/22 PKCα KO.

We transfected organotypic hippocampal slices from wildtype (WT) or PKCα knockout (KO) mice with small GTPase sensors and imaged CA1 pyramidal neurons using 2pFLIM. In response to glutamate uncaging targeted to a single dendritic spine (30 pulses at 0.5 Hz), the stimulated spine from WT slices rapidly enlarged by ∼150% (transient phase) and persisted with an increased volume of ∼40% lasting at least 25 min (Fig. 1b, c). This structural plasticity is associated with an increase in the functional strength of the stimulated spine (Gipson and Olive, 2017; Matsuzaki et al., 2004). Consistent with previous work, hippocampal CA1 neurons from PKCα knockout mice showed a deficit in sLTP regardless of sensor expression (Fig. 1c, (Colgan et al., 2018)). When sLTP was induced in single dendritic spines, we observed rapid and sustained activation of Rac1 (Fig. 1b, d) in the WT slices that was consistent with previously reported findings (Hedrick et al., 2016). However, we found that in the absence of PKCα Rac1 activation during sLTP is significantly attenuated (Fig. 1d). Rac1 activity in WT mice was restricted to the stimulated spines at early time points (2 – 4 min), but spread into the dendrite and nearby spines at later time points (10 – 20 min), consistent with a previous study (Hedrick et al., 2016). In PKCα KO mice, Rac1 activity both in stimulated spines and adjacent dendrites were inhibited (Fig. 1e). Thus, PKCα positively regulates Rac1 activation to facilitate plasticity.

In order to test the specificity of PKCα signaling toward Rac1, we tested whether the plasticity-induced activation of other small GTPases involved in plasticity, Ras or Cdc42, (Lai and Ip, 2013; Sit and Manser, 2011; Zhu et al., 2002), was also impaired in the absence of PKCα. Although the impairment of sLTP was observed in PKCα KO neurons transfected with Ras or Cdc42 sensor (Fig. 2a, c), the activation of these two small GTPases during sLTP were not affected by loss of PKCα (Fig, 2b, d). This demonstrates that Rac1 but not Ras or Cdc42 is downstream of PKCα during sLTP.

**Figure 2.**
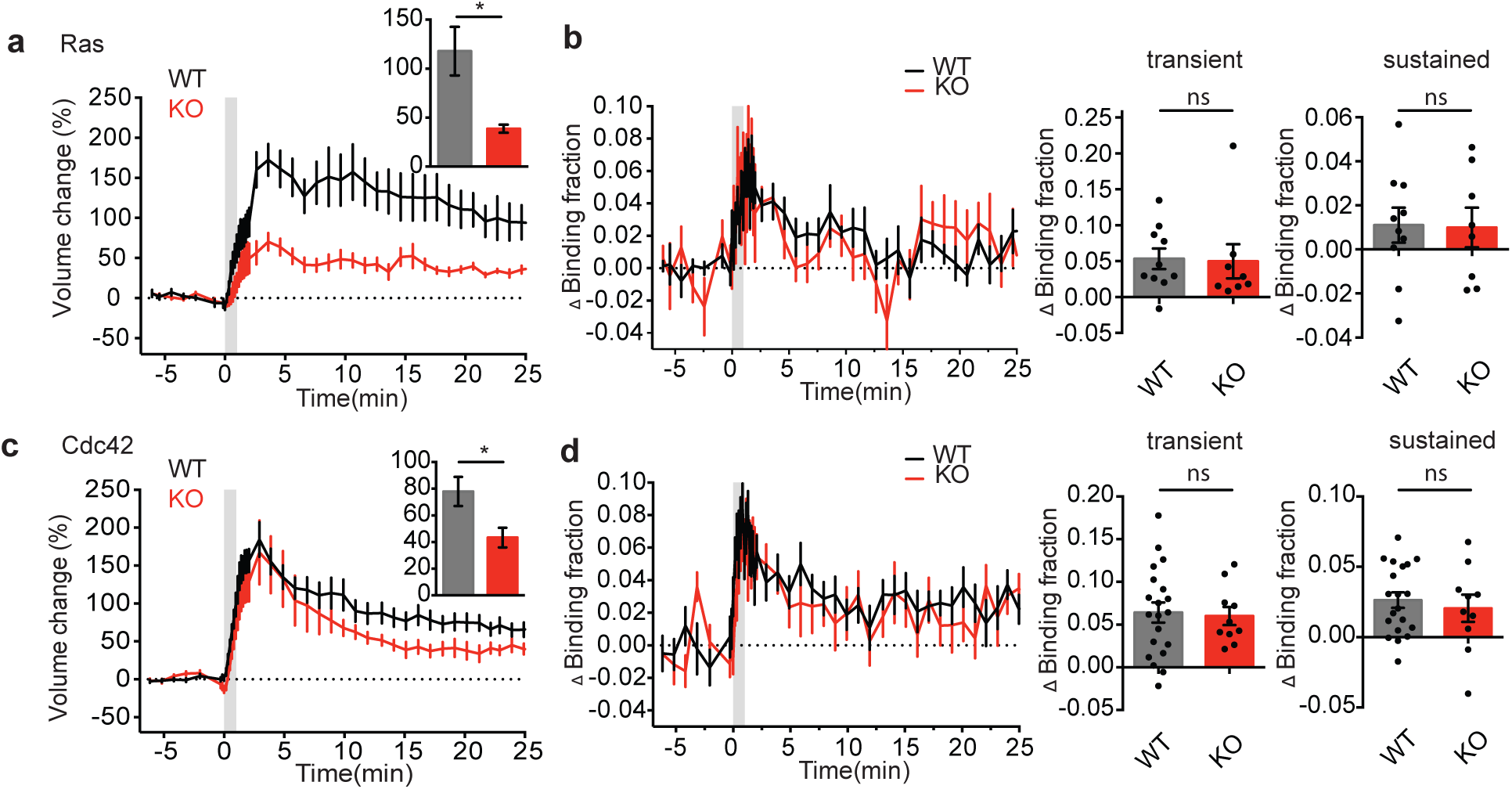
PKCα does not regulate Ras or Cdc42 activation during sLTP. **a**,**c**, Time courses and quantification of sustained (10–25 min) spine volume change induced by glutamate uncaging in neurons expressing Ras1 sensor (**a**) or Cdc42 sensor (**c**) from PKCα WT and KO littermates. **b**,**d**, Time courses and quantification of transient (1-3 min) and sustained (10-25 min) Ras1 (**b**) or Cdc42 (**d**) activation in stimulated spines from WT and PKCα KO littermates. Data are mean ± s.e.m. n(neurons/spines) Ras: n= 8/10 WT, n= 7/8 KO. Cdc42: n= 14/19 WT and n= 7/10 KO *P < 0.05, two-tailed t-test.

We next investigated whether the PDZ binding domain of PKCα, which was shown to be critical for its isozyme-specific signaling in spine plasticity (Colgan et al., 2018), was required for downstream regulation of Rac1 (Fig. 3a). PKCα or PKCα lacking its PDZ binding domain was sparsely and postsynaptically expressed alongside the Rac1 sensor in hippocampal slices from PKCα knockout mice. We found impairment of Rac1 activation and sLTP only when the PDZ binding domain was disrupted. This finding suggests a crucial role for PDZ binding domain of PKCα in Rac1 activation during sLTP (Fig. 3b).

**Figure 3.**
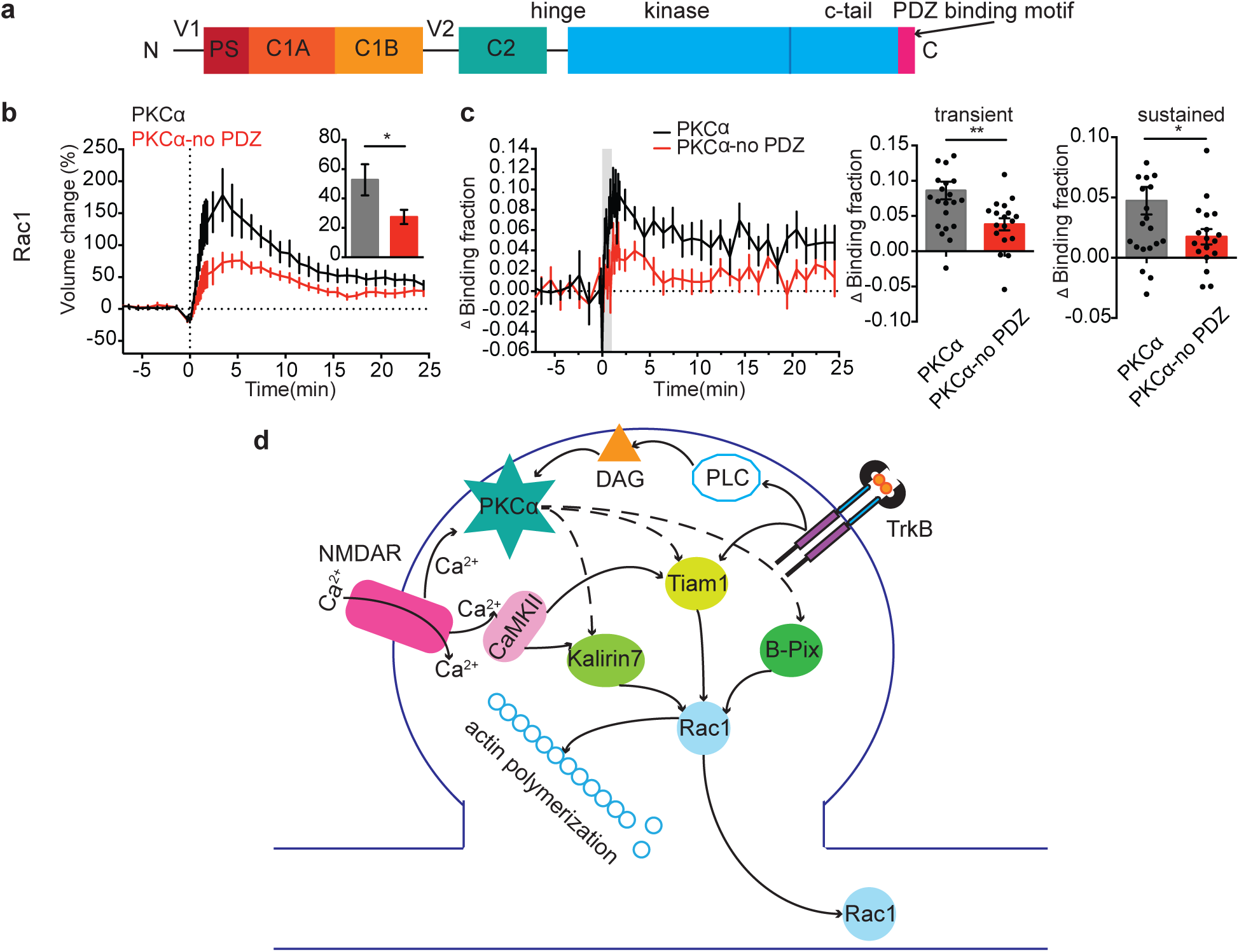
PKCα regulates Rac1 activation during sLTP via PDZ binding domain. **a**, Primary structure of PKCα showing pseudosubstrate, C1A and C1B domains, C2 domain, kinase domain, C-terminal tail, and PDZ binding motif. **b, c** Time courses and quantification of sustained (10–25 min) spine volume change **(b)** and transient and sustained Rac1 activation **(c)** induced by glutamate uncaging in PKCα KO hippocampal neurons expressing PKCα and PKCα without PDZ domain (PKCα-no PDZ). n = 20/23 PKCα and 14/18 PKCα-no PDZ (neurons/spines). Data are mean ± s.e.m. *P < 0.05, two-tailed t-test. **d**, Schematic of potential PKCα regulation of Rac1.

## Discussion

In this study, we have described that PKCα regulates the activation of Rac1, but not Ras or Cdc42, during sLTP of dendritic spines (Fig. 3d). This modulation relies on PKCα’s PDZ-binding motif, which may localize PKCα to a signaling complex scaffold that also recruits Rac1 GTPase-activating proteins (GAPs) or Guanine nucleotide-exchange factors (GEFs). Although the specific mechanism through which PKCα regulates Rac1 activity remains to be determined, potential scaffolds include PICK1, SAP-97 or PSD-95, which can interact with PKCα through its C-terminal PDZ binding domain and have been implicated in plasticity of spines (Callender and Newton, 2017; Ehrlich et al., 2007; Nakamura et al., 2011;; Staudinger et al., 1997; Volk et al., 2010; Waites et al., 2009).

The regulation of Rac1 during plasticity through GAPs and GEFs is highly complex in order to allow for tightly-regulated remodeling of various cytoskeletal domains in a precise spatial and temporal pattern. While PKCα could modulate Rac1 through inhibition of a GAP or Rho GDP dissociation inhibitors (RhoGDIs) (Garcia-Mata et al., 2011; Van Aelst and D’Souza-Schorey, 1997; Zhang et al., 1998), much of the specificity of Rac1 GTPase signaling is reportedly regulated by GEFs (Bellanger et al., 2000; Cook et al., 2014; Van Aelst and D’Souza-Schorey, 1997). At least ten different RhoGEFs have been identified to be localized to the post-synaptic density (Kiraly et al., 2010). Of these, Kalirin 7, Tiam1 and B-Pix all show preferential GEF activity for Rac1, are able to be phosphorylated by classic PKC isozymes in-vitro and are required for sLTP of spines (Buchanan et al., 2000; Chen et al., 2012; Fleming et al., 1999, 1997; Mandela and Ma, 2012; Mertens et al., 2003; Shirafuji et al., 2014; Saneyoshi et al., 2019). Kalirin7 and B-Pix also contain a PDZ binding domain which could localize them to PDZ containing-scaffolds together with activated PKCα (Park et al., 2003; Penzes et al., 2001; Saneyoshi et al., 2019). On the other hand, although Tiam1 itself contains a PDZ domain, the structure of its PDZ domain does not predict PKCα as a preferred binding partner (Shepherd et al., 2010).

Both Rac1 and PKCα activation are downstream of TrkB receptor activation in the stimulated spine (Colgan et al., 2018; Hedrick et al., 2016) (Fig. 3d). One important future direction is to determine if TrkB-dependent activation of Rac1 is solely through PKCα or whether a more complex feedback loop is in play. The increased activity and spreading of Rac1 promoted by PKCα is consistent with an essential role for PKCα in integrating neurotrophic signals to facilitate plasticity. Moreover, this link between PKCα and Rac1 supports a growing understanding of the molecular mechanisms underlying heterosynaptic facilitation, whereby the induction of plasticity in one spine lowers the threshold of plasticity induction in nearby spines (Colgan et al., 2018; Hedrick et al., 2016).

This study identified a novel molecular pathway that links short-lived calcium influx and activation of PKCα, to long-lasting Rac1 activation and changes in spine structure. Taken together with previous findings that another Ca^2+^-dependent kinase, CaMKII, also activates Rac1 during sLTP (Saneyoshi et al., 2019; Hedrick et al., 2016), Rac1 appears to be a key convergence point of multiple upstream calcium-dependent pathways (Fig. 3d). We anticipate that the better understanding of Rac1 signaling pathway may ultimately help us to uncover the signaling network with which memories are encoded.

## Materials and Methods

### Animals

All experimental procedures were approved by the Max Planck Florida Institute for Neuroscience Animal Care and Use Committee. P4-P8 mouse pups from both sexes were used for organotypic slices for imaging studies. The genotype of each animal was verified before preparing slices. PKCα KO 129/sv animals were received from Dr. Michael Leitges. Animals were crossed to C57Bl/6N Crl and are on a mixed background. For all the experiments WT littermates were used as controls for KO animals.

### Plasmids

All the plasmids used were previously developed and described in publication and are available on Addgene (Colgan et al., 2018; Hedrick et al., 2016; Oliveira and Yasuda, 2013). Briefly, the Rac1 sensor consisted of mEGFP-Rac1 (Addgene #83950) and mCherry-PAK2 binding domain-mCherry (Addgene #83951), the Ras sensor consisted of pCI-mEGFP-HRas ((Addgene #18666) and pCI-mRFP-RBD^K65E,K108A^-mRFP (Addgene #45149), the Cdc42 sensor consisted of mEGFP-Cdc42 (Addgene #29673) and mCherry-Pak3(60-113)/S74A/F84A-mCherry-C1 (Addgene #29676).

### Organotypic hippocampal slice cultures and transfection

Organotypic hippocampal slices were prepared from wildtype or transgenic postnatal 4-8 day old mouse pups of both sexes as previously described (Stoppini et al., 1991). In brief, the animals were anaesthetized with isoflurane, after which the animal was quickly decapitated and the brain removed. The hippocampi were dissected and cut into 350 µm thick coronal hippocampal slices using a McIlwain tissue chopper (Ted Pella, Inc) and plated on hydrophilic PTFE membranes (Millicell, Millipore) fed by culture medium containing MEM medium (Life Technologies), 20% horse serum, 1mM L-Glutamine, 1mM CaCl_2_, 2mM MgSO_4_, 12.9mM D-Glucose, 5.2mM NaHCO_3_, 30mM Hepes, 0.075% Ascorbic Acid, 1µg/ml Insulin. Slices were incubated at 37 °C in 5% CO^2^. After 7-12 days in culture, CA1 pyramidal neurons were transfected with biolistic gene transfer (O’Brien and Lummis, 2006) using 1.0 µm gold beads (8–12 mg) coated with plasmids containing 50 μg of total cDNA of interest in the following ratios. Rac1 sensor, donor: acceptor = 1:2; Rac1 sensor plus PKCα, donor: acceptor: PKCα = 1:2:1; Rac1 sensor plus PKCα without PDZ binding domain, donor: acceptor: PKCα-no PDZ domain = 1:2:1; Ras sensor, donor: acceptor = 1:3; Cdc42 sensor, donor: acceptor = 1:1). Neurons expressing all plasmid combinations were imaged 2–5 days after transfection.

### 2pFLIM

FLIM imaging using a custom-built two-photon fluorescence lifetime imaging microscope was performed as previously described (Murakoshi et al., 2011). 2pFLIM imaging was performed using a Ti-sapphire laser (Coherent, Cameleon) at a wavelength of 920 nm with a power of 1.4-1.6 mW. Fluorescence emission was collected using an immersion objective (60×, numerical aperture 0.9, Olympus), divided with a dichroic mirror (565 nm) and detected with two separated photoelectron multiplier tubes placed after wavelength filters (Chroma, 510/70-2p for green and 620/90-2p for red). Both red and green channels were fit with photoelectron multiplier tubes (PMT) having a low transfer time spread (H7422-40p; Hamamatsu) to allow for fluorescence lifetime imaging. Photon counting for fluorescence lifetime imaging was performed using a time-correlated single photon counting board (SPC-150; Becker and Hickl) and fluorescence images were acquired with PCI-6110 (National instrument) using modified ScanImage (Pologruto et al., 2003)(https://github.com/ryoheiyasuda/FLIMimage_Matlab_ScanImage). Intensity images for analysis of sLTP volume changes were collected by 128×128 pixels as a z stack of three slices with 1 µm separation and averaging 6 frames/slice. Spine volume was measured as the integrated fluorescent intensity of EGFP after subtracting background (F). Spine volume change was calculated by F/F_0_, in which F_0_ is the average spine intensity before stimulation.

### Two-photon glutamate uncaging

A second Ti-sapphire laser tuned at a wavelength of 720 nm was used to uncage 4-methoxy-7-nitroindolinyl-caged-l-glutamate (MNI-caged glutamate) in extracellular solution with a train of 4–8 ms, 2.8-3.0 mW pulses (30 times at 0.5 Hz) in a small region ∼0.5 µm from the spine of interest as previously described(Colgan et al., 2018). Experiments were performed in Mg^2+^ fee artificial cerebral spinal fluid (ACSF; 127 mM NaCl, 2.5 mM KCl, 4 mM CaCl_2_, 25 mM NaHCO_3_, 1.25 mM NaH_2_PO_4_ and 25 mM glucose) containing 1 µM tetrodotoxin (TTX) and 4 mM MNI-caged L-glutamate aerated with 95% O_2_ and 5% CO_2._ Experiments were performed at room temperature.

### 2pFLIM analysis

To measure the fraction of donor that was undergoing FRET with acceptor (Binding Fraction), we fit a fluorescence lifetime curve summing all pixels over a whole image with a double exponential function convolved with the Gaussian pulse response function:

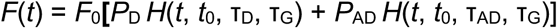

where τ_AD_ is the fluorescence lifetime of donor bound with acceptor, *P*_D_ and *P*_AD_ are the fraction of free donor and donor undergoing FRET with acceptor, respectively, and *H*(*t*) is a fluorescence lifetime curve with a single exponential function convolved with the Gaussian pulse response function:

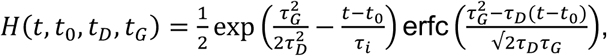

in which τ_D_ is the fluorescence lifetime of the free donor, τ_G_ is the width of the Gaussian pulse response function, *F*_0_ is the peak fluorescence before convolution and *t*_0_ is the time offset, and erfc is the complementary error function.

We fixed τ_D_ to the fluorescence lifetime obtained from free eGFP (2.6 ns), and then fixed τ_AD_ to fluorescence lifetime of the donor bound with acceptor (1.1 ns). For experimental data, we fixed τ_D_ and τ_AD_ to these values to obtain stable fitting. To generate the fluorescence lifetime image, we calculated the mean photon arrival time, <*t*>, in each pixel as:

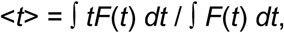

Then, the mean photon arrival time is related to the mean fluorescence lifetime, <*τ*>, by an offset arrival time, *t*_*o*_, which is obtained by fitting the whole image:

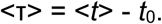

For small regions-of-interest (ROIs) in an image (spines or dendrites), we calculated the binding fraction (P_AD_) as:

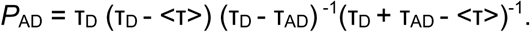

Data with lifetime fluctuations in the baseline that were greater than .15 ns were excluded before further analysis.

### Statistical analysis

All values are presented as mean ± SEM unless otherwise noted. Number of independent measurements (n[neurons/spines]) is indicated in figure legends. Unpaired two-tailed student’s t test was used for comparing two independent samples. Two-way ANOVA followed by multiple comparison test was used to compare grouped data sets (Prism 6, GraphPad). Data were only excluded if obvious signs of poor cellular health (for example, dendritic blebbing, spine collapse) were apparent.

## Acknowledgements

We would like to thank Dr. Michael Leitges for the PKCα KO 129/sv animals; M. Dowdy and the MPFI ARC for animal care; members of the Yasuda laboratory for discussion; L. Yan for helping with the microscope; and D. Kloetzer for managing the laboratory. This work was funded by NIH (R01MH080047, DP1NS096787) and the Max Planck Florida Institute for Neuroscience.

## Author details

### Xun Tu

- Neuronal Signal Transduction Group, Max Planck Florida Institute for Neuroscience, Jupiter, FL, USA.
- International Max Planck Research School for Brain and Behavior, Jupiter, FL, USA.
- FAU/Max Planck Florida Institute Joint Graduate Program in Integrative Biology and Neuroscience, Florida Atlantic University, Boca Raton, FL, USA.

## Contribution

Conceptualization, Designed the experiments, Data curation, Formal analysis, Validation, Investigation, Visualization, Writing—original draft, Writing—review and editing

## Ryohei Yasuda

Neuronal Signal Transduction Group, Max Planck Florida Institute for Neuroscience, Jupiter, FL, USA.

## Contribution

Conceptualization, Supervision, Designed the experiments, Writing—review and editing

## Contribution

Conceptualization, Investigation, Designed the experiments, Writing—review and editing

